# Patterned mechanical feedback establishes a global myosin gradient

**DOI:** 10.1101/2021.12.06.471321

**Authors:** Hannah J. Gustafson, Nikolas Claussen, Stefano De Renzis, Sebastian J. Streichan

**Affiliations:** Department of Physics, University of California Santa Barbara, Santa Barbara, California 93106, USA; Biomolecular Science and Engineering, University of California Santa Barbara, Santa Barbara, California 93106, USA; EMBL Heidelberg, Meyerhofstrasse 1, 69117 Heidelberg, Germany

## Abstract

Morphogenesis, the coordinated execution of developmental programs that shape embryos, raises many fundamental questions at the interface between physics and biology. In particular, how the dynamics of active cytoskeletal processes are coordinated across the surface of entire embryos to generate global cell flows is poorly understood. Two distinct regulatory principles have been identified: genetic programs and dynamic response to mechanical stimuli. Despite progress, disentangling these two contributions remains challenging. Here, we combine *in toto* light sheet microscopy with genetic and optogenetic perturbations of tissue mechanics to examine theoretically predicted dynamic recruitment of non-muscle myosin II to cell junctions during *Drosophila* embryogenesis. We find dynamic recruitment has a long-range impact on global myosin configuration, and the rate of junction deformation sets the rate of myosin recruitment. Mathematical modeling and high frequency analysis reveal myosin fluctuations on junctions around a mean value set by mechanical feedback. Our model accounts for the early establishment of the global myosin pattern at 80% fidelity. Taken together our results indicate spatially modulated mechanical feedback as a key regulatory input in the establishment of long-range gradients of cytoskeletal configurations and global tissue flow patterns.

---

Active regulation of the cytoskeleton represents an important set of cell scale processes capable of driving morphogenesis in a variety of contexts(*1–3*). Just as unbalanced forces in a tug of war lead to motion, tissue scale gradients of subcellular localized cytoskeletal components create force imbalances that quantitatively predict tissue flows (*4–7*). To understand morphogenesis, then, requires a quantitative understanding of how such gradients are created. In some cases, relations between gene expression patterns establishing the body axis and the distribution of cytoskeletal components have been uncovered by molecular investigations (*8, 9*). For example, the anisotropic myosin distribution driving germ-band extension (GBE) in *Drosophila* depends on Toll family receptors downstream of pair rule genes (*10*). Although recent work suggests a link between Tolls and Rho activity upstream of myosin, the quantitative picture of the molecular pathway that converts Toll patterns to anisotropic myosin localization remains incomplete (*2, 11*).

In addition to genetic inputs, growing evidence indicates that mechanical cues can influence the cell properties and behaviors that determine how a tissue deforms (*12–15*), a class of phenomena we here refer to as “mechanical feedback”. While a feedback loop via Rho activation has been indicated in cell culture (*16, 17*), molecular analysis suggest that this mechanism might not be present in all organisms (*18*). Other studies report local myosin recruitment in response to mechanical deformation, however, the molecular mechanism *in vivo* remains unknown (*12*). Here, the role played by tissue-specific gene expression in controlling mechanical cues complicates the task of disentangling the impact of a pure genetic deterministic program from the effects of dynamic, post-translational mechanical feedback mechanisms (*2*). Furthermore, mechanical coupling of cells affects how forces are transmitted across tissues (*19, 20*) to activate putative mechanosensitive processes (*21–23*). Due to lack of tools for characterizing mechanics during organismal development (*24*), it remains unclear what inputs cells sense, how far these effects reach, and what relevance they may have on morphogenetic outputs.

In this work we take an integrated approach, combining quantitative investigation with theory and quantitative analysis. Noll et al (*13*) developed a physical theory of tissue mechanics proposing “dynamic recruitment” of myosin as a mechanism to ensure mechanical stability of a tension dominated network. In its simplest form, dynamic recruitment predicts a specific quantitative signature, according to which rates of myosin are recruited proportional to rates of cell edge deformation. We test this hypothesis by combining light sheet microscopy (*25*), optogenetics (*26*), and quantitative analysis to characterize myosin-II dynamics on adherens junctions in response to the deformation of cell edges. As an experimental system we focused on germband extension (GBE) during *Drosophila* gastrulation. GBE is characterized by the convergent extension of cells which leads to more than two-fold elongation of the body axis, and it involves cells immediately adjacent to the ventral furrow VF (*8*). This is a major tissue deformation process driven by the collective contraction and invagination of approximately 1,000 cells from the ventral surface of the embryo (*27*). The remaining epithelial surface maintains mechanical integrity, such that cells outside of the VF become stretched to compensate for the greatly reduced number of cells spanning the same circumference (*27*). These highly dynamic and overlapping processes result in substantial tissue flows and represent an ideal context to study the quantitative nature of dynamic recruitment. Our results demonstrate that cytoskeletal dynamics can be captured in a physical model that quantitatively accounts for the establishment of myosin-II patterns from strain rates and indicate that long range gradients of cytoskeletal and tissue flow patterns are determined by spatially patterned mechanical feedback.

## Results

### Myosin rate and strain rate are correlated and graded along the DV axis

To quantitatively test whether cells measure edge deformation and proportionally recruit myosin (Fig 1A), we use *in toto* live imaging of fluorescently labeled non-muscle myosin II (myosin) at sub-cellular resolution with confocal multi view light sheet microscopy(*25, 28*). We first characterized the dynamics of myosin in wild type embryos during VF formation and the fast phase of GBE by measuring junctional myosin normalized to the cytoplasmic pool (junctional accumulation). Measuring myosin in this way allows for quantification of rates of changes in local density caused by myosin motors binding to the actin cortex at adherens junctions independent of the concentration of fluorescently marked motors (see SI for more detail). In a region of the germ-band close to the VF, myosin junctional accumulation increased significantly over the course of VF formation, specifically on junctions parallel to the dorsoventral (DV) axis (Fig 1B). When this same measurement was applied to the entire surface of the embryo, a clear gradient with high myosin junctional accumulation adjacent to the VF and little to no junctional myosin on the dorsal region became evident (Fig 1C). This gradient along the DV axis suggests that myosin dynamics vary by region. We thus divided the trunk into discrete regions along the DV axis and measured the myosin accumulation averaged within these regions as a function of time (Fig 1D,E). Myosin accumulation in the region closest to the VF was initially low at the onset of VF formation and increased linearly during invagination, to then decrease after furrow internalization was completed (Fig 1D). When analyzing myosin accumulation over time by region, we found that the rate of accumulation was spatially graded, steepening the gradient in myosin levels over the course of VF formation (Fig 1E).

**Figure 1:**
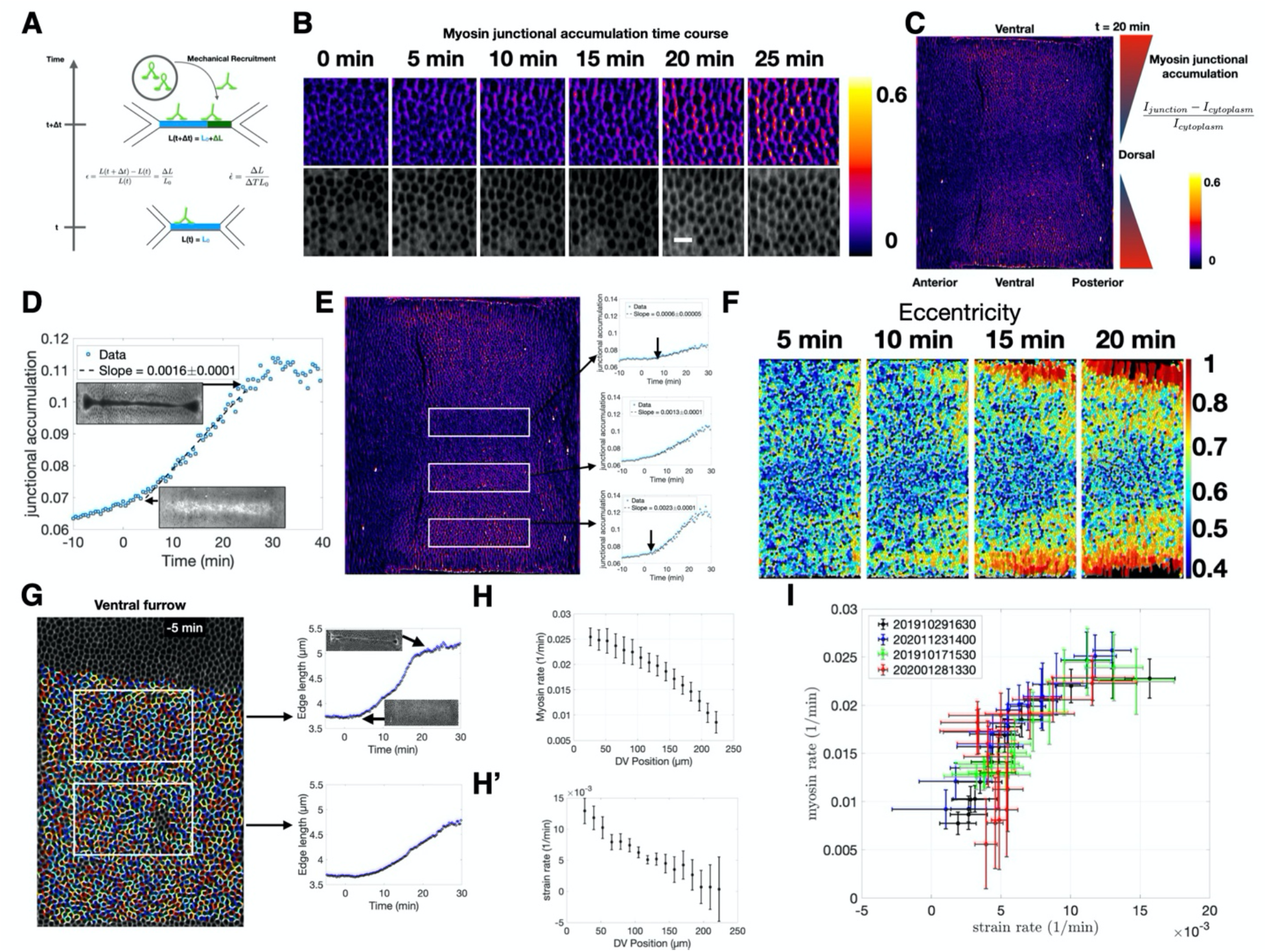
Correlation between the rate of strain and the rate of myosin recruitment on individual junctions. **(A)** Schematic representation of the proposed mechanical feedback mechanism. Equations give definitions of strain and strain rate. **(B)** Junctional myosin accumulation over time (see C for definition) in a region of the lateral ectoderm of a sqh::mCherry expressing embryo. t = 0 corresponds to 10 minutes before the onset of ventral furrow formation. Bottom row cell surface outlines 4 μm below adherens junction. **(C)** Junctional myosin accumulation across the entire surface of an embryo during early germ-band elongation (20 minutes after ventral furrow formation begins). Color gradient in triangles represents graded junctional myosin accumulation along D-V axis. **(D)** Junctional accumulation over time on cell edges located in a single region of the embryo (corresponding to the middle box in E). Insets show morphological landmarks in the region of the VF in sqh::mCherry expressing embryo at the corresponding time point. **(E)** From the same movie as in (C). White boxes indicate regions for which the plots of junctional myosin accumulation over time are measured. Arrows indicate the earliest time point through which the line fitting the data passes. **(F)** Cell eccentricity over time in a region corresponding to the central area of the germ-band. Pullback is as in (C), except for regions anterior to the cephalic furrow which are not shown **(G)** Example of cell segmentation and single edge tracking used to measure strain rate over time in the two regions designated by the white boxes. Insets show the same morphological landmarks as D. **(H)** Myosin rate measured as a function of position along the DV axis. (H’) Strain rate measured as a function of position along the DV axis. **(I)** Plot of strain rate vs. myosin rate for four control embryos. Each cross corresponds to the measurements as shown in H and H’. All error bars are SEM unless otherwise indicated. For detailed discussion of number of cell edges per embryo, see SI.

We next characterized the strain of cell edges during this time frame. Cells nearest the VF were stretched towards the ventral midline giving rise to an anisotropic cell shape oriented towards the VF (Fig 1F). While this effect was first observed in cells immediately adjacent to the furrow, it rapidly spread across the DV axis as furrow formation progressed. Single cell edge tracking revealed cell edges oriented parallel to the DV axis and adjacent to the VF were strongly strained (i.e. elongated) as VF forms (SI movie 1). This effect decreased systematically with increasing distance from the furrow (Fig 1G). Therefore, both the rate of myosin recruitment (Fig 1H) and the rate of strain (Fig 1H’) showed a clear trend with the highest rates immediately dorsal to the VF and steadily decreasing along the DV axis. Plotting the myosin accumulation rate versus the strain rate revealed a strong correlation between the two (Fig 1I). While the relation between genetic patterning and myosin anisotropy during GBE is well documented, it is currently unknown how a DV myosin gradient emerges on cell edges (*9, 29*). Our analysis demonstrates that the characteristic anisotropic myosin pattern of the germband occurs concomitant with VF formation and that the germband begins to move as the DV myosin gradient steepens. These observations suggest that the strain generated by the VF plays a role in establishing the global myosin anisotropy pattern during GBE.

### Strain rate induces myosin recruitment

Testing whether a causal mechanism underlies the relation between strain and myosin recruitment rates requires non-invasive methods capable of modifying strain rates (*26, 30*). We turned to optogenetics to transiently activate the cytoskeleton in spatially restricted domains to induce cell contraction and monitored the resulting changes in cellular flow, strain, and myosin dynamics in adjacent regions (Fig 2A, Fig S3A-D’). To this end, we transiently activated a Cry2-CIBN based RhoGEF2 optogenetic module with a spatially patterned infrared beam from a femtosecond laser (*26*). As described previously(*31*), this protocol results in two-photon optogenetic activation, allowing for spatially restricted photo-activation patterns to control regional contractility (see methods, SI). We utilize this strategy to induce strains in various regions and directions and observe the response of myosin.

**Figure 2:**
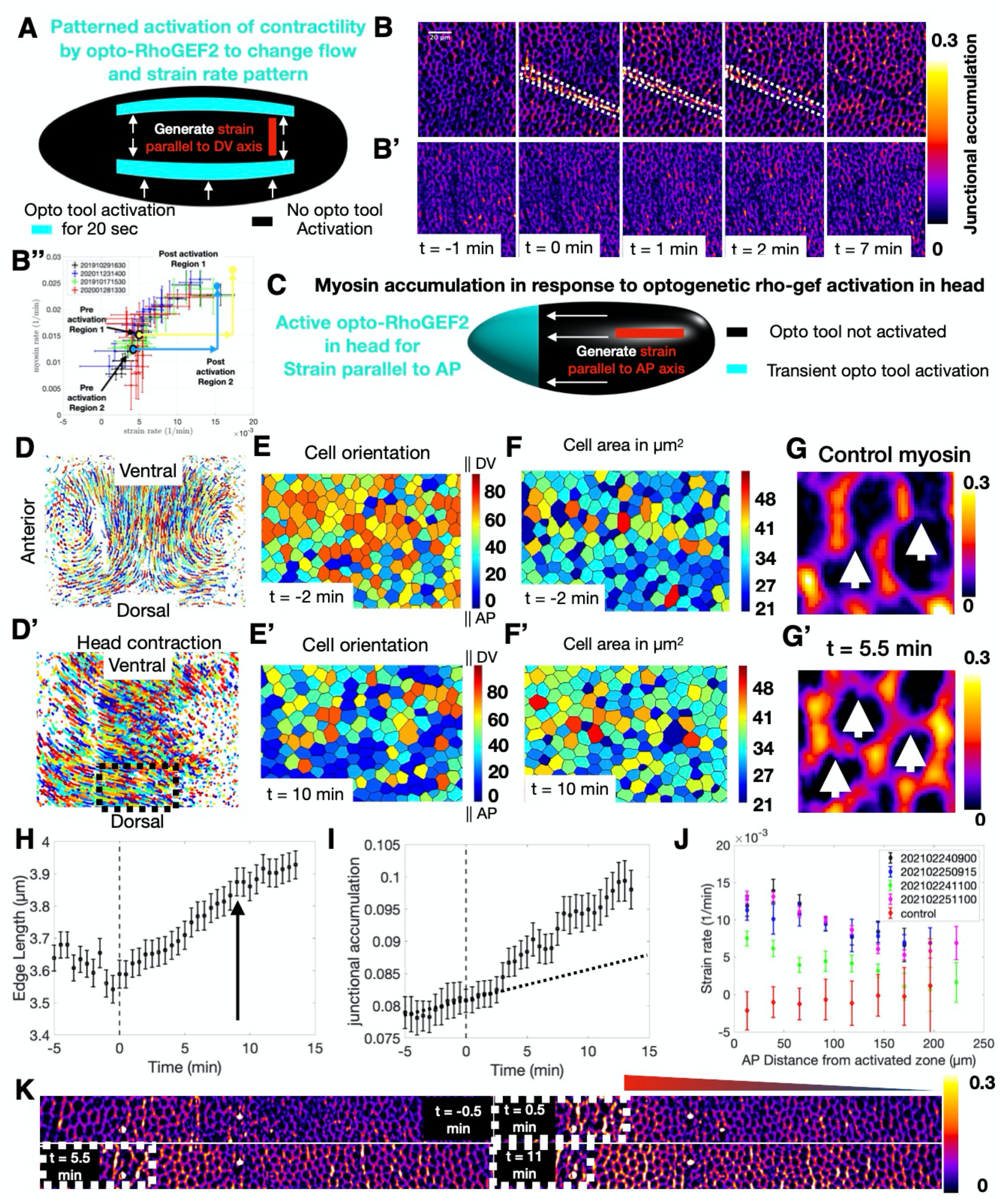
Patterned contractility by optogenetics redirects flows. **(A)** Schematic of the optogenetic strategy employed to generate strain parallel to the DV axis. Cyan area indicates the area of transient optogenetic activation. Opto-RhoGEF2 is not activated in black regions. White arrows depict direction of resulting flow. Red bar indicates axis of strain. **(B)** Junctional myosin accumulation in one of the experiments depicted in A. t=0 corresponds to the first image taken after activation. Domain of activation is contained in white dashed box. **(B’)** Junctional myosin accumulation in a control embryo in an equivalent region and developmental stage as the activated embryo shown in B. **(B’’)** Myosin and strain rates in optogenetically activated embryos follow the same relation observed in control during VF formation (Fig 1I). Myosin and strain rate of two realizations of the experiment depicted in (A) before and after optogenetic activation. **(C)** Schematic of optogenetic strategy employed to generate strain in the AP direction. Cyan area indicates the area of transient optogenetic activation in the head. Opto-RhoGEF2 is not activated in black regions. White arrows again represent direction of induced flow. Red bar indicates axis of strain. **(D,D’)** Cell trajectories on one lateral side of a control embryo (D) compared to the cell trajectories in an activated embryo (D’) at the same developmental time. **(E,E’)** Cells in the region of the black dashed box in (D’) color coded by orientation measured as the angle of the cell long axis relative to the AP axis before (E) and after (E’) activation. **(F, F’)** Same region as E,E’ color coded for apical cell area before (F) and after(F’) activation. **(G)** Junctional myosin accumulation in a control embryo. **(G’)** Junctional myosin accumulation in an equivalent region and developmental time as (G) in an activated embryo. White arrows identify junctions parallel to the AP axis, and direction of strain with significantly higher myosin accumulation compared to corresponding junctions in a control embryo. **(H)** Junction length over time of junctions parallel to the AP axis in an activated embryo. Gray dashed line shows time of activation. Black arrow indicates the time point at which the rate of junction length change begins to decrease. **(I)** Accumulation of myosin on the same junctions as H. Gray dashed line shows time of activation. Black dashed line shows the myosin accumulation expected without activation based on the slope of the pre-activation data points. **(J)** Strain rate on junctions parallel to AP axis as a function of distance from activation zone along the AP axis for four different head activated embryos. Red points show no significant deformations parallel to AP axis in control embryos. **(K)** Myosin junctional accumulation over time along the AP axis in a lateral region of a head activation experiment. White dashed line shows boundary between infrared activated (left) and unactivated (right) cells. Triangle shows gradient corresponding to graded junctional myosin accumulation along the AP axis.

As shown above, during VF formation the endogenous patterns of both strain and myosin rates are anisotropic, with much higher values on edges parallel to the DV-axis than their orthogonal counter parts. This raises the question of whether mechanical feedback itself is anisotropic. If it is not, the direction of strain has the capacity to affect myosin anisotropy. To distinguish between these two possibilities, we induced strain of cells either along the DV or AP direction. First, we stretched cells along the DV direction, focusing our analysis on the dorsal region. We induced photo-activation in two parallel regions, extending across the AP axis (Fig 2A). The first timepoint after activation (t = 0 min) showed a strong signal in the myosin channel, confined to the illumination pattern, resulting from RhoGEF2 recruitment to the membrane and Rho signaling stimulation (Fig 2B Fig S4A-A’’). Importantly, in the first 30 seconds following activation, cells intervening the two infrared illumination patterns did not show any detectable changes in junctional myosin levels, while myosin signals in the activated regions strongly increased (Fig S4A,A’). Therefore, we conclude that optogenetic activation remained confined to the infrared illumination pattern and did not activate surrounding cells. Within 1 minute, cells in the activated regions contracted (Fig 2B, Fig S4A’), and caused a modification of cell flow pattern in non-activated regions (Fig S4B,B’). While cells remained nearly stationary immediately prior to activation, all cells in the intervening region flowed towards the activation lines and stretched following activation (Fig S4A-B’). About 2 minutes after activation, we observe a pronounced increase in junctional myosin within this region oriented preferentially along the direction of strain (Fig 2B, Fig S4A’’). In contrast, cells in the same region and developmental time in control embryos showed low accumulation of junctional myosin (Fig 2B’). Quantitative analysis revealed a significant change in cell shape (Fig S4C,C’), concomitant with increased myosin recruitment rates to the junction (Fig S4D,E). Remarkably, the increase of strain rate was matched by a proportional increase in myosin rate, similar to the relation observed in the whole embryo analysis of VF formation (Fig 2B’’). However, during VF formation similarly high strain and myosin accumulation rates were only observed in close vicinity of the VF but not on the dorsal surface (Fig 1G, H).

Next, we designed an activation scheme to generate deformations along the AP axis (Fig 2C, Fig S3E-H’). Because inducing strain in this direction requires overpowering the underlying flow pattern, we transiently activated RhoGEF2 in the entire head region with the aim to generate strain of comparable level to that produced during VF invagination along the DV direction. Optogenetic activation resulted in nearly uniform head contraction, producing a striking reorientation of cell flow (Fig 2D,D’). In control embryos the primary flow pattern, as established by single cell tracking, was dominated by cells of the lateral ectoderm flowing towards the ventral midline (Fig 2D). Head activation caused a global reorientation of flow with cell trajectories pointing towards the head (Fig 2D’), while retaining a component in the ventral direction due to VF formation. To characterize the effects of this altered flow profile on cell geometry, we quantified cell orientation and area before and after activation. While initially cells were oriented along the DV direction (Fig 2E), within 10 minutes of activation cells on the dorsal side were reoriented along the AP axis, consistent with the pulling direction (Fig 2E’). Cell area showed a modest, but significant increase after activation (Fig 2F,F’). Therefore, this activation scheme reorients both cell flow and geometry along the direction of the imposed ectopic strain. It is well established that during GBE cell interfaces parallel to the AP direction do not accumulate myosin (*32*). Indeed, in control embryos these junctions accumulate very little myosin, such that they were hardly visible relative to perpendicular junctions (Fig 2G). In contrast, high levels of myosin accumulation on these junctions could be observed in head activated embryos (Fig 2G’).

Quantitative analysis confirmed that junctions parallel to the ectopic strain direction extend significantly within 1 min after activation and continued to grow at a steady rate for about 8-10 minutes (Fig 2H). The rate of myosin accumulation on these junctions was initially very low and similar to that measured on the dorsal surface of control embryos (Fig 1E, top panel) but increased within two to three minutes following activation (Fig 2I). These results demonstrate a strict temporal sequence of events: (I) accumulation of myosin restricted to the photo-activation domain (II) contraction of activated cells initiates a change in flow within one minute and alterations of strain rate patterns on edges (III) proportionally increased myosin accumulation on strained edges two to three minutes after activation.

Next, we measured the strain rate as a function of distance from the activation front. Adjacent to the activation site, the perturbation drastically increased the strain rate, which gradually decayed towards the posterior pole (Fig 2J). Depending on the strength of activation, the propagating effects of cell reorientation could still be distinguished for up to 200 μm, corresponding to roughly half the length of the embryo (Fig 2J). Consistent with this observation, the level of myosin accumulation showed a clear gradient that was higher close to the region of activation and decreased towards the posterior end (Fig 2K). This gradient in the AP direction recapitulated that observed during VF formation, only rotated 90 degrees in line with the direction of the ectopic strain. In addition to this AP pattern, we also noted that VF invagination induced flow towards the ventral region causing stretching of junctions parallel to the DV axis (Fig 2D’). Consistently, we also observed myosin accumulation on junctions parallel to the DV axis characteristic of this stage.

Taken together, these results indicate that strain rate quantitatively affects the pattern of myosin recruitment on both DV and AP junctions. Specifically, the orientation of a junction with respect to the direction of the strain determines the level of junction deformation and, thereby, the level of myosin recruitment. Our results further uncover the long-range consequence of local mechanical perturbations: strain can be transmitted and sensed hundreds of microns away from the source, causing a deformation rate that leads to a proportional increase in the rate of myosin accumulation.

### Strength of feedback depends on DV position but not edge orientation

Our experimental strategy allowing monitoring the response of every junction on the surface to mechanical deformations, prompted us to study mechanical feedback in deeper quantitative detail. The well-established link between cytoskeletal activity and gene expression patterns (*9, 33*), raises the possibility that genetic patterning regulates mechanical feedback. Such an effect might become visible by studying the proportionality coefficient between strain rate and myosin, which we refer to as feedback coefficient, as a function of position across the surface of the embryo. A non-uniform coefficient would suggest that the strength of mechanical feedback, and thus the amount of myosin recruited to a given deformation, is patterned.

To explore this possibility, we turn to a more detailed quantitative analysis comparing the response to deformations on the dorsal vs ventral surface. We created strain parallel to the DV axis (Fig 2A) at varying DV positions and found that in all cases strain and myosin rates increased in response to perturbation (Fig S4G,H). Strikingly, while strain rate showed no correlation with DV position, myosin rate was systematically lower at the dorsal vs. ventral surface (Fig S4G,H). This suggests the ratio of myosin vs strain rate, and thereby the mechanical feedback coefficient relating these two quantities, is graded along the DV axis.

We created strain parallel to AP axis as described before (Fig 2C), and despite nearly uniform contraction induced along the circumference, we observed striking differences at the dorsal vs. ventral regions, both in terms of cell trajectories (Fig 3A,A’) and cell shapes (Fig 3B,B’), as well as myosin activation patterns (Fig 3C,C’). Quantitative analysis focusing only on junctions parallel to the AP axis confirms that junctions on the dorsal surface experience higher strain compared to their counter parts in ventral regions (Fig 3D). Notably, myosin rate did not exhibit the same trend, and in fact the average myosin rate was lower on the dorsal than on the ventral region (Fig 3E). Analysis of myosin accumulation rate as a function of strain rate for junctions parallel to the AP axis revealed a lower feedback coefficient in dorsal than ventral regions (Fig 3F,F’).

**Figure 3:**
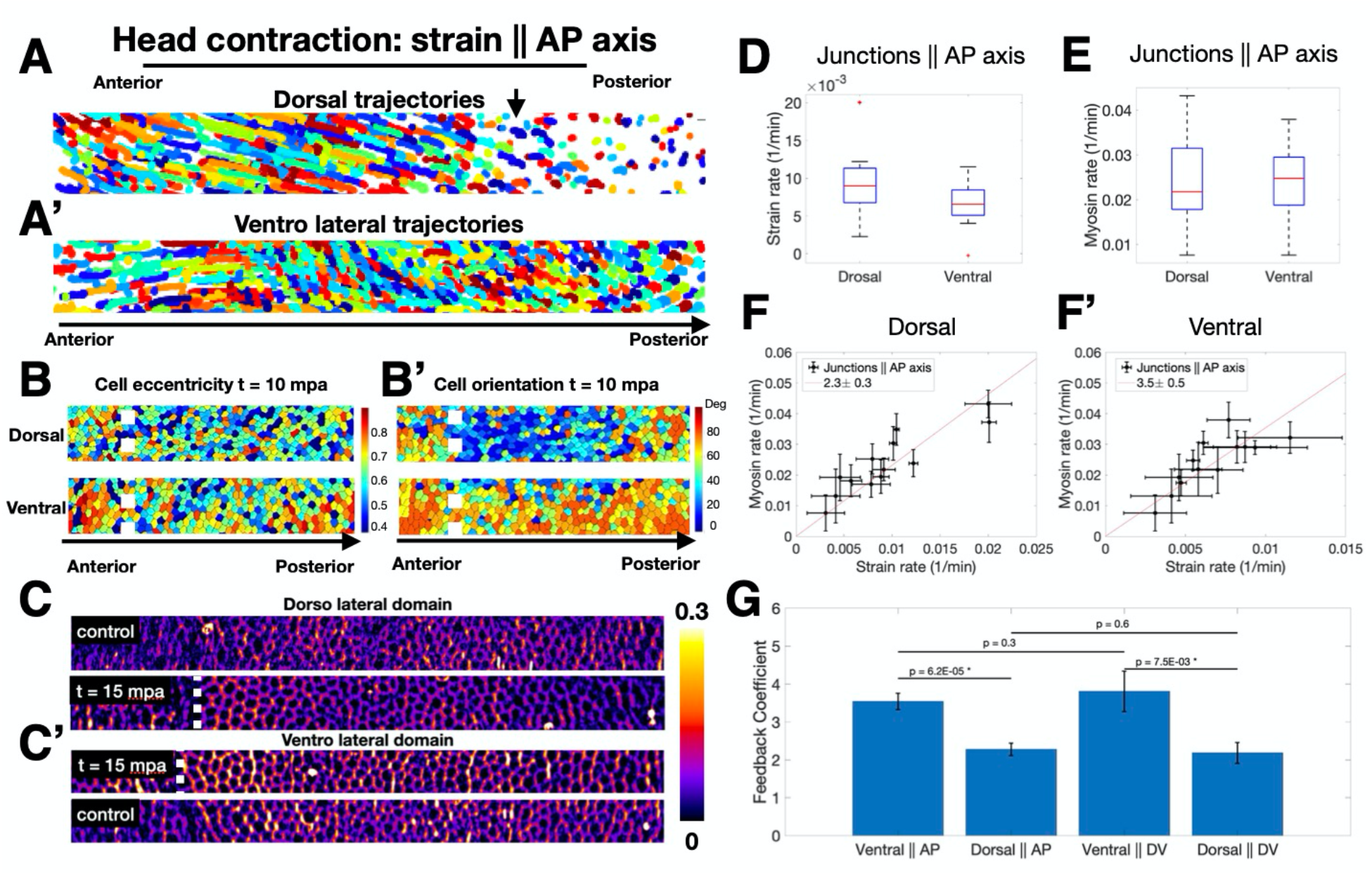
Regional analysis of mechanical feedback coefficient and directionality. **(A,A’)** Trajectories of cells tracked for 20 minutes following head activation in dorsal (A) and ventrolateral (A’) regions of the embryo approximately 50 um wide along the AP axis. **(B)** Same regions as (A) and (A’) 10 minutes after activation color coded for eccentricity. **(B’)** Same regions as (B) color coded for orientation of the cell long axis relative to the AP axis. **(C)** Junctional myosin accumulation in the dorsolateral region of control (top) and activated (bottom) embryos at the same developmental stage. **(C’)** Junctional myosin accumulation in the ventrolateral region of activated (top) and control (bottom) embryos at the same developmental stage. Regions in C,C’ are 30 um wide spanning the AP axis. **(B-C’)** White dashed line shows boundary of activation region in the anterior. **(D)** Strain rate in junctions parallel to the AP axis in dorsal and ventral regions following head activation. **(E)** Myosin rate on junctions parallel to the AP axis in dorsal and ventral regions. **(D-E)** Red line indicates median, box lower and upper quartiles, whiskers are minimum and maximum. **(F,F’)** Strain rate vs myosin rate on junctions parallel to the AP axis in the dorsal (F) and ventral (F’) regions for all head activation experiments, N=5 embryos. Red line shows best fit proportionality of the data points. Legend shows feedback coefficient and the 95% confidence interval. Error bars are SEM. **(G)** Comparison of feedback coefficient based on junction orientation (parallel to DV vs AP) and region (dorsal vs ventral). This coefficient is the proportionality between strain rate vs myosin rate plot as in (F,F’), Fig S4GH. P values are obtained by single sided Welch t-test.

Fig 3G shows a summary of the feedback coefficient quantified as the ratio of myosin rate over strain rate, organized in the four different categories we analyzed: dorsal vs ventral domains, and junctions parallel to AP vs DV axis. These results do not show a significant difference in terms of junction orientation within either dorsal or ventral region but do across regions. This indicates that mechanical feedback acts independently of junction direction, and that genetically encoded spatial patterning of the mechanical feedback coefficient downstream of position information determines the amount of myosin recruitment in response to a given deformation.

### Single junction analysis reveals two timescales govern myosin dynamics

The preceding analyses average cell junction strain and myosin rates within local regions of interest and rely on data at 30s temporal resolution. While this is sufficient to identify causal relationship between strain and myosin rates, we wanted to analyze the relation of deformation and myosin at high frequency. We moved to local imaging at higher temporal resolution using confocal microscopy to track both junction length and myosin signal over time. Representative images (Fig 4A) show a junction marked with both myosin and a membrane marker at successive time points. Single edge tracking revealed that myosin density on the edge (Fig 4B) and junction length (Fig 4B’) exhibited rich dynamics with rapid fluctuations around a longer time scale trend. From myosin concentration and junction length, we extracted strain and myosin rates and plotted them as a function of time (Fig 4C). Myosin and strain rates oscillated out of phase with a period of 74 seconds (Fig 4D). Cross correlation between myosin and the strain rates showed a negative correlation with a time shift of −7 secs and a positive correlation with a time shift of 30 secs, suggesting close to half a period phase shift. This effect is specific to the relationship between myosin and strain rates as cross-correlation between strain rate and the concentration of the membrane marker GAP43::mCherry showed no significant peaks.

**Figure 4:**
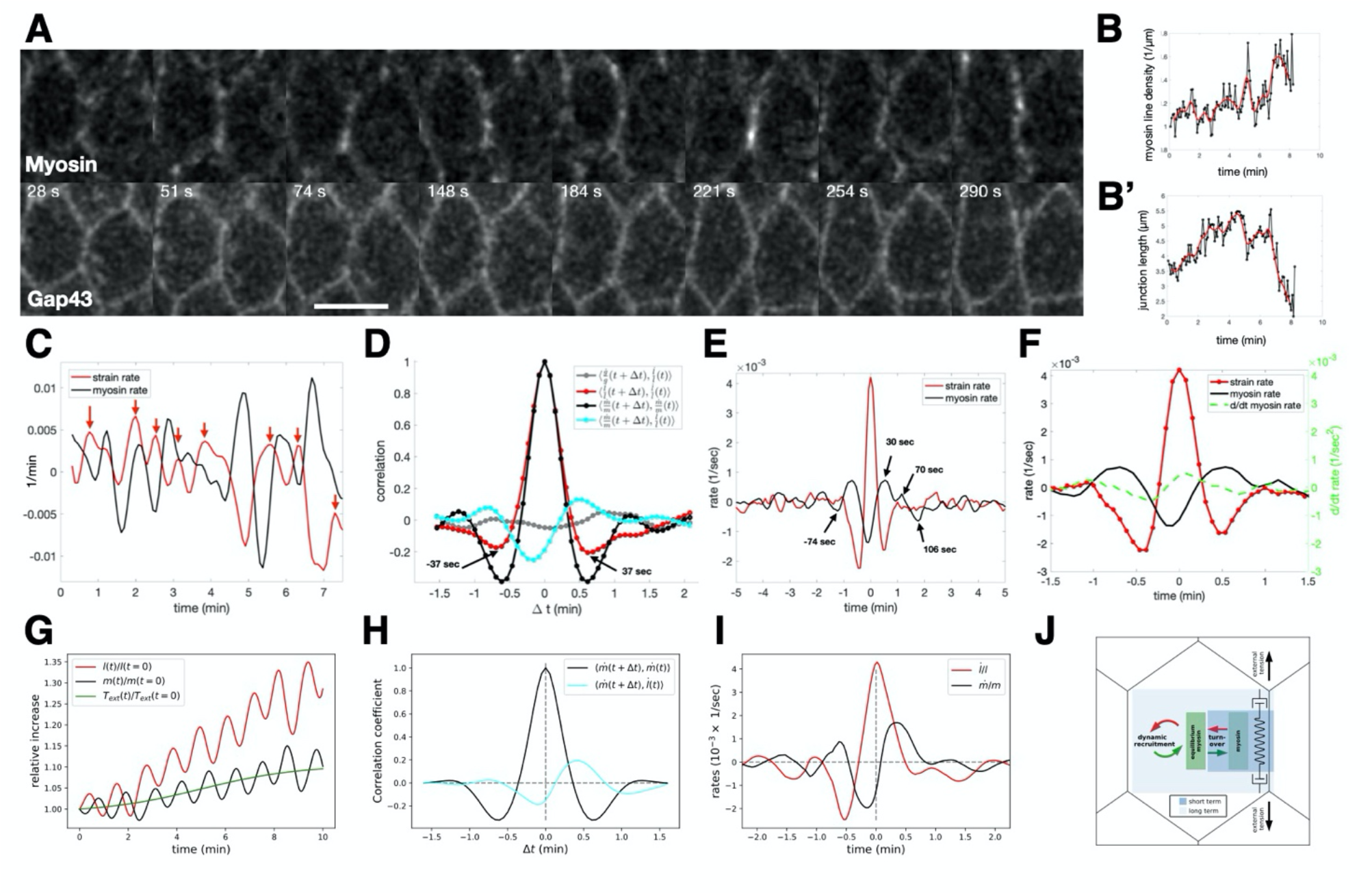
Single edge analysis and modeling of myosin and membrane dynamics. **(A)** Confocal images of cells during early gastrulation marked with sqh::GFP and Gap43::mCherry showing timepoints that highlight the oscillations of myosin and junction length. Scale bar is 5 microns. **(B)** Myosin line density normalized to initial myosin line density over time. **(B’)** Junction length in microns over time. In both (B,B’) black points represent data. The Red line is the fit through the data points. **(C)** Plot of strain rate (red line) and myosin rate (black line) over time calculated from the data plotted in (B,B’). Red arrows indicate peaks in strain rate. **(D)** Autocorrelation analysis of strain rate (red) and myosin rate (black) and cross-correlation analysis of membrane rate with strain rate (gray) and myosin rate with strain rate (green). **(E)** Traces obtained by aligning strain rate peaks and averaging the aligned strain rate (red) and myosin rate (black). **(F)** Same as in (E), with displayed data corresponding only to one cycle before and after t = 0 and with the myosin acceleration plotted on the same graph (green dashed line). **(G)** Sample simulation of the relative increase in junction length (red) and myosin concentration (black) over time given by the model. External tension applied to junction shown in green. **(H)** Autocorrelation analysis of myosin rate (black) and cross-correlation of myosin rate with strain rate (green), based on N=500 model simulation runs. **(I)** Average curves of strain rate (red) and myosin rate (black) predicted by the model after strain rate peak alignment, as in E (N=500). (J) Cartoon representation of the dilution oscillator model with mechanical feedback.

To characterize the longer term impact of strain rate on myosin on the same junction, we investigated the typical response of myosin rate to significant peaks in the strain rate (red arrows in Fig 4C) by averaging the time traces of multiple edges centered around peaks (Fig 4E,F). We focused on the fluctuating forces acting on tracked junctions to investigate the consequences of peaks in the strain rate. The average strain rate around the peak has, as expected, a global maximum at 0 and two adjacent minima symmetrically localized around plus/minus 30 seconds. The average myosin rate exhibited a global minimum shortly before the strain rate peaks, as expected from the cross-correlation analysis. The peaks of myosin appeared symmetrical both in timing as well as magnitude. The situation was different for the minima. While the last minimum in myosin rate appeared 74 seconds before the peak, the first minimum after the peak did not appear until 106 seconds. This suggests a prolonged phase of myosin sourcing in response to deformation. Indeed, the time derivative of the myosin rate became positive as soon as the junction extended, signified by a positive strain rate (Fig 4F). Taken together, these findings indicate that myosin is subject to two separate time scales: rapid fluctuations around a slowly changing reference value set by mechanical-driven recruitment, similar to theoretical ideas proposed in (*34*).

To quantitatively explore this hypothesis, we formulated a physical model with few parameters. We adapted a “concentration oscillator” model, which has been shown to capture the dynamics of apical myosin and cell area fluctuations (*35*), by incorporating a mechanical feedback mechanism, which was proposed as a requirement to maintain mechanical stability of tissues (*13*), see SI for more detail. With this model, we could capture both the short-term oscillations observed in the single edge dynamics as well as the longterm drift in junction rest length and myosin levels (Fig 4G). We assumed that at short timescales, a cell edge behaves as an elastic spring that can elongate under external tension or shorten, based on the balance of external forces, elasticity, and actomyosin contractility. Oscillations between these two states are sustained by the effects of dilution, concentration, and myosin turnover, which are captured by the concentration oscillator. Simulating the autocorrelation and cross-correlation analysis using this model accurately recreated the measures above (Fig 4H). At longer timescales, the cell edge undergoes viscoelastic relaxation due to cytoskeletal remodeling, allowing for changes in the junction rest length. Mechanical feedback causes recruitment of myosin in proportion to the edge strain which balances the external tension to stabilize junction length, leading to plastic deformation and adjusted equilibrium myosin levels (Fig 4G). By modeling these behaviors, we were able to recreate the peak aligned strain rate and myosin rate curves, including key features such as the phase-shifted oscillations and asymmetry about zero in both curves (Fig 4I). Notably, a concentration oscillator model alone, without mechanical feedback, was insufficient to account for this observation (see SI Fig 5). In summary, external tension on a junction influences myosin dynamics at two timescales: fast turnover characterized by oscillatory behavior without net gain of myosin, and slower adjustment of myosin levels due to mechanical feedback (Fig 4J, see SI for more detail). Taken together, the high frequency analysis of single junctions and the accuracy with which our model reproduces these dynamics supports the mechanism for dynamic myosin recruitment identified through our analysis of wild type and optogenetic whole embryo live imaging data.

**Figure 5:**
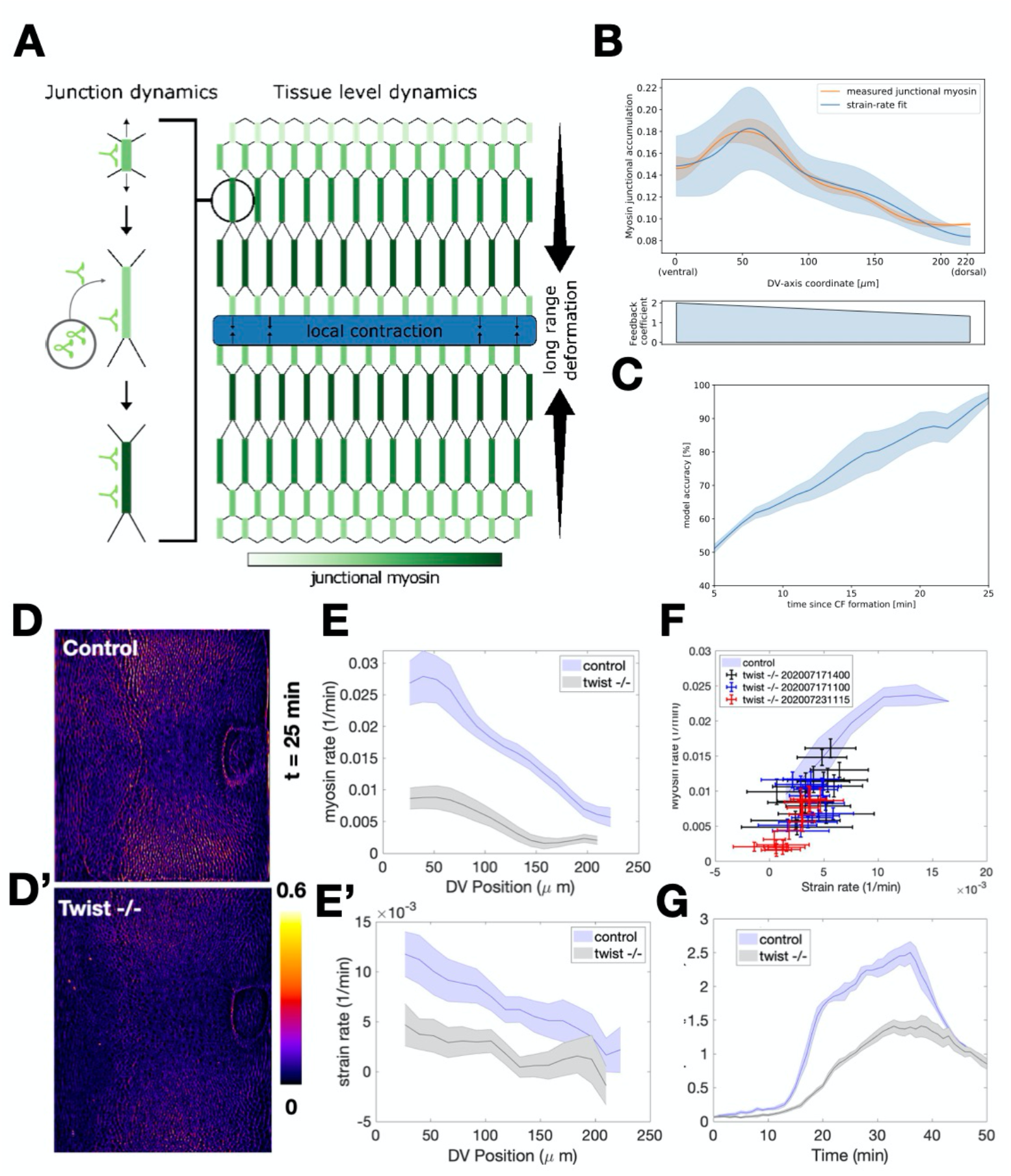
Prediction of embryo-scale myosin distribution from junctional model and analysis of twist mutant embryos. **(A)** Illustration connecting junctional dynamics (left) with tissue-scale dynamics (right). Local contraction propagates across the tissue as long-range deformations, which serve as the source of external tension activating mechanical feedback for myosin recruitment and establishing tissue scale gradients of junctional myosin accumulation. **(B)** Comparison of measured junctional myosin levels (orange line) and myosin levels predicted from model using total strain as input (blue line). Lower plot shows DV gradient of the feedback coefficient input into the model. **(C)** Model accuracy over time, representing level of agreement between predicted and measured myosin profiles. **(D-D’)** Junctional myosin accumulation in twist heterozygous (‘control’, D) and twist homozygous (‘twist’, D’) embryos at equivalent times. **(E)** Myosin rate as a function of D-V position in control (blue) and twist (black) embryos. **(E’)** Strain rate as a function of D-V position for control (blue line) and twist (black) embryos. **(F)** Strain rate vs myosin rate plot with the average wild type curve (gray) and individual twist embryos (points). **(G)** Mean velocity over time for wild type and twist mutant embryos. t = 0 corresponds to the onset of CF formation.

### Embryo scale myosin levels can be accurately predicted from integrated strain rate

Next, we wanted to test if our single junction model could be used to account for the early myosin dynamic patterns established during VF formation at the whole embryo scale. Because myosin accumulation rate is likely influenced by factors other than strain rate, it remains unclear if the predictive power of mechanical feedback will extend to the large scale and long-time pattern of myosin dynamics, when the anisotropic pattern characteristic of GBE is established. i.e. during the 20 minutes following cephalic furrow formation. By integrating the deformation rates of individual junctions, we used our model to provide a quantitative estimate of the contribution from mechanical feedback to the time course of myosin on junctions. As shown in Fig 1C myosin pattern exhibited a striking gradient along the DV axis. On the scale of single cell edges, our model predicts that a junction elongating under external tension will undergo a net increase in myosin levels due to mechanical feedback (Fig 5A, left). Local contraction in a mechanically coupled epithelium will generate external tension on junctions surrounding the site of contraction (Fig 5A, right). This effect decays with distance from the source of contraction, resulting in a gradient of external tension. Junctions with orthogonal orientation to the contraction pattern will thus experience an unbalanced external tension, extend, and recruit myosin (Fig 5A, right).

We used single edge tracking to reconstruct the history of deformations experienced by individual edges across the DV axis (Fig S2A-B). The myosin profile at any given time was quantified using the junctional accumulation measure (Fig 1C). Our model relates myosin dynamics to deformations by the feedback coefficient, which varies across the surface (Fig 3I). Therefore, we measured all spatially varying observables relevant to predicting the myosin profile from deformation rates, leaving only a single fitted constant in the integrated feedback law (see Theory SI for more detail). The predicted profile and the measured myosin were remarkably similar (Fig 5B). Agreement between our prediction and the observed myosin pattern is initially low, due to the overall low strain and myosin levels at early time points, but rapidly increased to ~80% just before the onset of GBE when myosin accumulation is highest, (Fig. 5C). This suggests our model is highly accurate both at the level of individual junctions, and across the entire DV axis.

This level of predictive power indicates that during GBE myosin distribution strongly depends on strain pattern. Because VF invagination is a major source of deformation in the early embryo, it is expected to contribute significantly to the generation of strain. This can be tested in mutants affecting VF formation, such as loss of function alleles of the transcription factor Twist. *twist* mutant embryos still exhibit residual cell behaviors that are hallmarks of VF formation. The high resolution of our analysis pipeline at the whole embryo level allowed us to exploit this detail in a quantitative test of our model. In *twist* mutants, Snail expressing cells in the ventral region undergo pulsed apical contractions, without intervening stabilization of the apical surface (*33*). This suggests these cells can still pull on the adjacent germband, albeit less effectively than their WT counterparts. We therefore expect a quantitatively reduced profile of deformations and accordingly predict a reduced myosin profile. Indeed, the levels of junctional myosin accumulation during early GBE were significantly reduced in *twist* mutants compared to wild type embryos at an equivalent developmental stage (Fig 5D,D’). Both myosin rates (Fig 5E) and strain rates (Fig 5E’) were 2.5 times lower and displayed shallower gradients in *twist* mutants. When plotting myosin rates vs strain rates, the *twist* data followed the same curve as their WT counterparts, although at lower strain and myosin rates (Fig 5F) indicating that the mechanisms underlying this proportionality were preserved in *twist* mutants. The mechanical consequences of the reduced myosin rate, and hence reduced myosin DV gradient, were apparent when comparing the mean velocity profile in wild type and *twist* embryos. This analysis revealed that the magnitude of flow during GBE, particularly during the fast phase, was also reduced by the same factor of 2.5 (Fig 5G). Despite the lower magnitude of flow, the direction of the flow pattern is largely maintained in *twist* embryos, such that GBE proceeds, albeit markedly slower. This remarkable quantitative agreement strongly suggests that deformation rates underlie the establishment of the early DV myosin gradient which sets the pace of germband extension.

## Discussion

Here we combined light sheet microscopy with physiological deformations afforded by patterned optogenetic contractility, to demonstrate a mechanical feedback mechanism which results in dynamic myosin recruitment to junctions proportional to the rate of edge deformations. Our data show that mechanical feedback is isotropic and patterned along the DV axis. High frequency analysis of individual cell junctions combined with a physical model revealed the dynamic nature of myosin on adherens junctions: similar to medial pools on the apical and basal cell surface, the junctional myosin concentration oscillates around a mean value. We show that the dynamics of this mean value can be quantitatively captured by a mechanical feedback mechanism which responds to the rate of edge strain. Our model also quantitatively bridges the junctional scale to the global embryo-scale, predicting the establishment of *in toto* myosin profiles from measured deformations. This level of predictive power is surprising, as genetic evidence in fixed samples demonstrates that pair rule genes affect myosin anisotropy via a recently discovered toll receptor code(*9*), yet we have not explicitly accounted for any A-P patterning in our model.

Our analysis of myosin dynamics further reveals that the strength of mechanical feedback is globally graded and patterned. This suggests that genetic patterning provides positional information to set the stage for mechanical feedback capacity of individual cells. In the context of gastrulation, these results suggest that patterning of the feedback coefficient along the DV axis acts as a developmentally planned physical mechanism for symmetry breaking, ensuring robust and globally graded myosin distribution. Reducing the DV gradient of the myosin profile reduces the speed of GBE accordingly.

Our optogenetic perturbations argue that the cytoskeleton plays a permissive role in the establishment of mechanical feedback in all directions, indicating that the molecular components of this mechanism are likely uniformly distributed at the cell surface. Intriguingly, Toll receptors are distributed uniformly (*10*), raising the possibility that mechanical feedback bridges the discrepancy in the current model between isotropic Toll distribution and anisotropic myosin distribution downstream of Tolls. In this scenario, strain anisotropy leads to anisotropic activation of an isotropic mechanical feedback pathway, potentially involving Toll receptors, to establish the anisotropic myosin distribution.

Future studies combining dynamic analysis with genetic perturbations of the pair rule genes and their targets may help reaching a unified quantitative model of cytoskeletal dynamics during *Drosophila* gastrulation. The highly conserved morphogenetic processes of apical constriction and cell intercalation driving VF formation and GBE (*36, 37*), respectively, raise the possibility that the mechanism we describe here may have implications for the development of a wide variety of species employing these modules. It is therefore intriguing to note a dual role for mechanics in development: not only as the set of physical properties that determine how tissues deform, but also as a genetically encoded physical mechanism to ensure robust tissue flow.

## Materials and methods

### Light sheet microscopy

We used a custom built Multi View Selective Plain Illumination Microscope (MuVI SPIM) (*28*) in a scatter reducing imaging mode (*25*) for fluorescent based live imaging of full *Drosophila* embryos at subcellular resolution. Briefly, the setup involves a pair of orthogonally arranged illumination and detection arms, in duplicate arranged such that illumination and detection face one another. For illumination we used a custom-built laser combiner that housed a set of continuous wave laser lines (488 nm, 561 nm, 660 nm) all OBIS LX, Coherent Inc.. These laser lines where always operating at the same set output power for each experiment. Stability of the beam was assessed using an optical power meter from Thorlabs (PMD 100D with S121C). These laser lines were combined using dichroic mirrors on kinematic mirror mounts. A broad band beam splitter from Omega Optical Inc., designed to evenly split the beam at 488 nm, 561 nm, and 940 nm (for use in conjunction with optogenetic activation, see below), was used to duplicate the light path for feeding the two illumination arms. The light paths in each illumination arm were identical. They first consisted of a pair of kinematic mirrors for alignment purposes, a galvanometric mirror (Cambridge technology), a scan lens (Sill Optics), a Tube lens (200 mm focal length) and a water-dipping objective (CFI Plan Fluor 10x, NA 0.3, Nikon). The detection involves a water dipping objective (APO LWD 25x, NA 1.1, Nikon), a filter wheel (Lambda, Sutter Instruments) with emission filters (FF01-542/27-25, FF01-609/62-25,BLP01-568R-25, BLP01-664R-25, all Semrock), tube lens (200 mm focal length), and an sCMOS camera (Hamamatsu ORCA-Flash 4.0 V3).

For optimal image quality we reduced optical scattering, following the strategy outlined in (*25*). Briefly, we used a National Instruments Multi-function I/O card (PCI-6229) to generate a sequence of electronic signals to synchronize galvanometric mirror phase with the start of image acquisition, and sample motion. The camera was operated light-sheet mode readout, at maximum speed setting for the acquisition front. The width of exposing pixels around this from was set to 52 pixels (corresponding to effective 13.4 μm) in all experiments.

Optical sectioning involved electronically controlled stages all from Physik Instrumente GmbH and Co.Kg. A translation stage (linear piezo stage P-629.1cd with E-753 controller), a rotational piezo stage (U-628.03 with C-867 controller), and a linear actuator (M-231.17 with C-863 controller).

Electronic acquisition was controlled through micro manager (*38*).

### Optogenetic activation

For tight spatial confined optogenetic activation we took advantage of the two photon effect described for fluorescence microscopy in (*39*). This non-linear method uses a high intensity infrared laser to allow for spatially restricted activation of a fluorophore or optogenetic constructs and is widely used in deep tissue imaging, e.g. neuro science (*26, 40*). Briefly, specificity is achieved because excitation of the chromophore requires simultaneous absorption of two infrared photons instead of one photon of ~half the wavelength normally employed. The energies required for a non-zero cross-section of this event are only achieved in the focal spot of the objective. This effect therefore drastically reduces light scattering thus avoiding optogenetic activation away from the focal plane of the objective. We used a tuneable femtosecond laser (Chameleon Vision II laser system, Coherent Inc.), set to 940 nm for spatially controlled activation

### Image acquisition

Embryos were dechorionated following standard procedures, and mounted in agarose gels as previously described(*28*). Imaging proceeds as follows: the embryo is imaged simultaneously from two objectives with sections spaced 1.5 um aprt, producing two separate Z stacks from opposite sides of the embryo. The embryo is rotated by 45 degrees and imaged again for 3 additional positions, for a total of 8 views per time point. Total imaging duration per time point is approximately 18 seconds allowing for temporal resolution of 30-60 secs.

### Data fusion

Fluorescent beads (Fluoresbrite multifluorescent 0.5-μm beads 24054, Polysciences Inc.) are diluted 1:1,000 in 1% low-melting point agarose solution into which the embryo is mounted for imaging. These fluorescent beads serve as fiducial markers to register views using interest point detection and matching with the Fiji plugin Multiview Reconstruction (*41*). The Difference of Gaussians interest point detection method is used to identify the bead locations in all views for all timepoints. Beads are matched using the fast 3D geometric hashing (rotation invariant) algorithm and all-to-all time point matching (global optimization). The views are then registered using an affine transformation model regularized to a rigid model with a lambda value of 0.10. Images are fused using Multiview deconvolution with an efficient Bayesian iteration. The PSF estimation is extracted from the beads. The resulting image has an isotropic resolution of 0.2619 um.

### Surface of Interest extraction

Tissue cartography was used to define SOI and extract fluorescence data on pullbacks (*42*). The embryo surface is determined using the Ilastik detector, giving rise to point cloud for SOI construction. This point cloud is fitted by using the sphere-like fitter class, to create a smooth representation of the SOI using cylinder coordinates defined by AP and DV axes.

### Confocal imaging

Embryos were dechorionated mounted on a glass bottom MaTek dish, and covered with water. Embryos were imaged with a Leica SP8 confocal microscope (HC PL Apo CS2 40x, NA 1.1, Water, Leica Inc.) An image stack corresponding to the apical surface (9 stacks with 1um axial resolution, 0.08 um lateral resolution) was acquired every 4.6-4.8 seconds during the time just before ventral furrow formation through the fast phase of GBE.

### Optogenetics crosses

Crosses between UASp>CIBN::pmGFP; UASp>Cry2::RhoGEF2 virgins and sqh::mCherry/Cyo; osk>Gal4/TM3 males were set up and kept in the dark at 25°C. All subsequent procedures were preformed using only red-light illumination. F1 progeny of the appropriate genotype were sorted and maintained in an embryo collecting cage. See Supplementary Information for more details.

### Segmentation and cell edge tracking

For segmentation of cell outlines, a layer approximately 5 μm below the SOI was segmented using Ilastik (*43*). The image mask generated from this workflow allows for edge detection and tracking using procedures outlined in(*44*), implemented in a custom matlab script, from which edge length and myosin intensity are measured. All measurements involve the metric tensor to correct for possible distortions from projections in maps (*42*). See SI for more detail.

### Fly lines

*halo twi^ey53^*/Cyo,sqh::GFP

*halo sna^IIG05^*/Cyo,sqh::GFP

w[*]; GAP43:: mCherry(attp40)/Cyo,sqh::GFP

UASp>pmGFP:CIBN, UASp>Cry2::RhoGef2

w[*], P[w+, UASp>CIBN]; P[w+, UASp>RhoGEF2-CRY2]

w[*]; P[w+, sqh::mCherry]/Cyo; P[w+, Osk>Gal4::VP16]

## Notes

### Competing Interest Statement

The authors have declared no competing interest.

